# Higher-order dissimilarity in biodiversity: Identifying dissimilarities of spatial or temporal dissimilarity structures

**DOI:** 10.1101/2022.11.22.517446

**Authors:** Ryosuke Nakadai, Keita Fukasawa, Taku Kadoya, Fumiko Ishihama

## Abstract

1. Elucidating biodiversity patterns and their background processes is critical in biodiversity science. Dissimilarity, which is calculated based on multivariate biological quantities, is a major component of biodiversity. As spatial and temporal biodiversity information availability increases, the scope of dissimilarity studies has been expanded to cover various levels and types of spatio-temporal biodiversity facets (e.g., gene, community, and ecosystem function), and diverse pairwise dissimilarity indices have been developed. However, further development of the dissimilarity concept is required in comparative studies on spatio-temporal structures of biodiversity compositional patterns, such as those exploring commonalities of biogeographical boundaries among taxa, compared to the conventional ones to consider higher dimensions of dissimilarity: dissimilarity of dissimilarity structures.
2. This study proposes a novel and general concept, high-order dissimilarity (HOD), for quantitatively evaluating the dissimilarities of spatial or temporal dissimilarity structures among different datasets, proposes specific implementations of HOD as operational indices, and illustrates the potential resolution of scientific and practical questions through HOD.
3. We further demonstrate the advantages of the HOD concept by applying it to actual patterns, such as long-term and/or large-spatial hypothetical monitoring datasets.
4. Our conceptual framework on HOD extends the existing framework of biodiversity science and is versatile, with many potential applications in acquiring more valuable information from ever-increasing biodiversity data.

## Introduction

Nature’s patterns are ubiquitous regardless of the level of biological organisation, and the elucidation of their determinants has been a long-standing and fundamental issue in biodiversity science. Dissimilarity is a key element of such patterns and has been studied in terms of spatial and temporal differences and changes in various biological entities, mainly at the same level of biological organisation (Anderson et al. 2011). This encompasses differences and changes in genetic alleles, species composition, interaction networks, and ecosystem functions (Whittaker 1960, 1972; Raymond & Rousset 1995; Mori et al. 2015; Poisot et al. 2012). The scope of dissimilarity is continuously expanding, covering various levels and types of biodiversity facets, which has stimulated the development of conceptual and analytical frameworks for the analysis of the determinants of dissimilarity patterns using various dissimilarity indices (Koleff et al. 2003; Lozupone & Knight 2005; Baselga 2010; Rocchini et al. 2018; Mammola & Cardoso 2020). Notably, despite the varieties, all these indices have a standard form in that one dissimilarity value is calculated from two vectors of multivariate biological quantities (i.e., pairwise dissimilarity indices), and the calculation of dissimilarity values for all pairs of vectors yields a dissimilarity matrix (Anderson et al. 2011).

Some of the spatio-temporal structures of biodiversity composition patterns, however, are beyond the scope of pairwise dissimilarity and identifying, quantifying, and understanding them necessitates higher-order considerations. In a spatial context, higher-order considerations are useful for the analysis of categorically different dissimilarity matrices, such as among assemblages of different taxonomic groups and among species (i.e., comparative biogeography and phylogeography). For example, biogeographical researchers have attempted to identify common biogeographical borders among taxa (Wallace 1894; Holt et al. 2013; Whittaker et al. 2013; Komaki 2021); similarities or dissimilarities in spatial composition dissimilarities among multiple taxa need to be evaluated. In the temporal context, higher-order consideration would provide a framework for quantitatively evaluating dissimilarity in compositional changes (i.e., temporal asynchrony of compositional changes), such as among sites and different trophic groups. For example, temporal changes in community composition (i.e., temporal beta diversity: Legendre 2019; Nakadai 2022) have recently been expanded to multiple sites at large spatial scales (Magurran et al. 2019; Nakadai 2020; Gotelli et al. 2022), thus necessitating quantitative evaluation of inter-site dissimilarity between temporal dynamics of community composition measured using pairwise comparison. Furthermore, loss and biotic homogenisation in species composition and intraspecific haplotypes have been studied in-depth since identifying human-induced impacts on biodiversity within the context of conservation (Olden 2006; Valtonen et al. 2017). Moreover, quantifying the temporal change of spatial dissimilarity structure from a baseline at a time point is essential to assessing the impacts. Despite the apparent importance and scientific need for higher-order consideration in dissimilarities, general concepts and formal statistical methods for quantitative comparisons across conventional pairwise dissimilarities have not been fully developed.

To address this limitation, we introduce a general framework for considering dissimilarities at a higher order (hereafter referred to as higher-order dissimilarity; HOD) than that of conventional pairwise dissimilarities. HOD is a natural extension of the pairwise dissimilarity indices and considers differences between dissimilarity matrices. In addition, it can be a novel building block in the studies of the dissimilarities of spatio-temporal structures of biodiversity compositional patterns. First, in the present study, we formally define the novel and general concept of HOD to quantitatively evaluate dissimilarities of spatial or temporal dissimilarity structures among different datasets (Fig. 1a). Second, we designed the concept of HOD and proposed its implementation as a general statistical method. Third, we demonstrated the advantages of the HOD concept by applying it to actual patterns, such as long-term and/or large-spatial hypothetical monitoring datasets. Finally, we discuss the potential impacts of the concept of “higher-order dissimilarity” and the developed analytical framework on a wide range of research fields, including understanding complex spatio-temporal structures in biodiversity and future methodological challenges.

**Figure 1.**
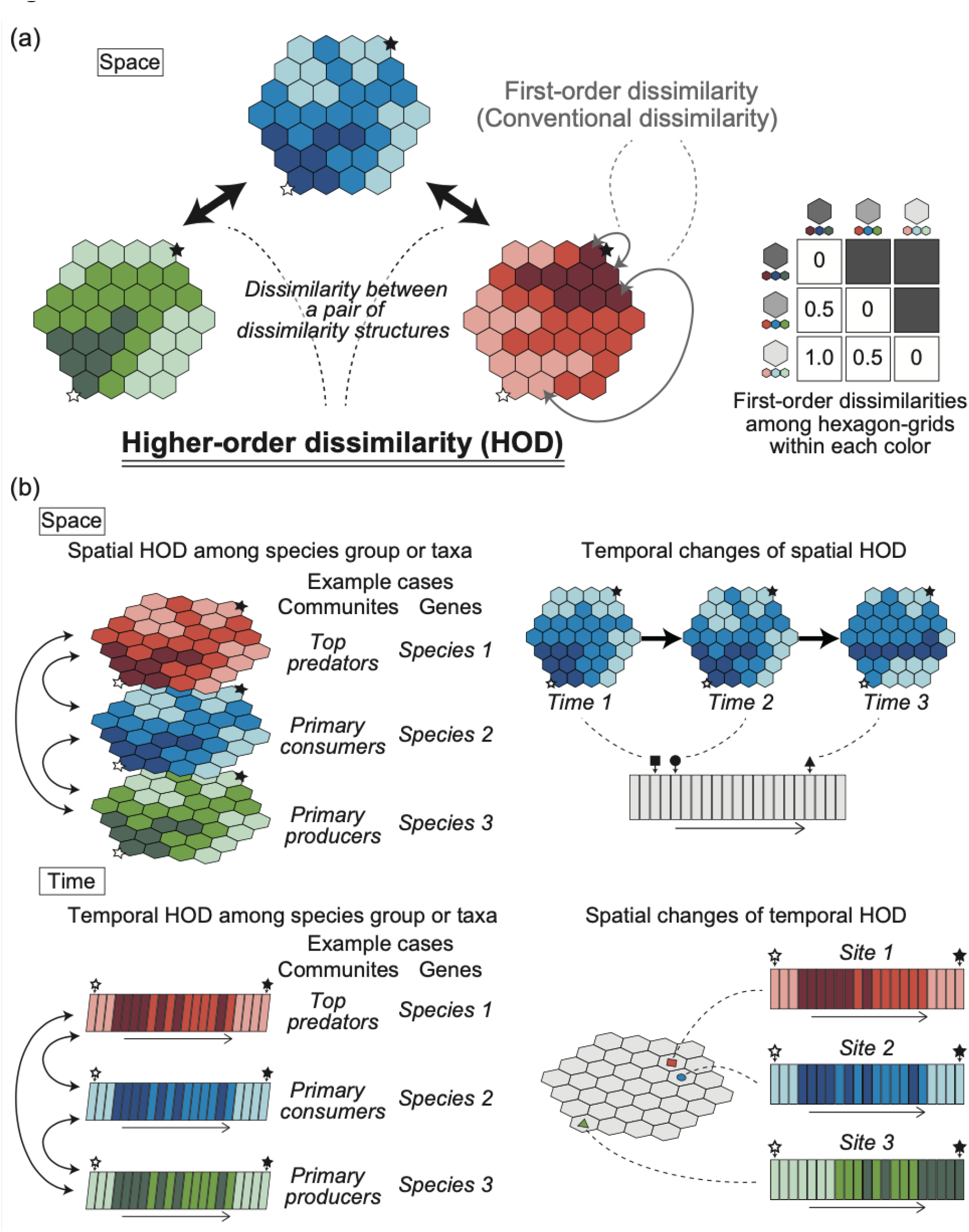
Schematic of higher-order dissimilarity (HOD). A spatial case of HOD is shown in (a). Although conventional dissimilarity has focused on dissimilarity among the same dimension, higher-order dissimilarity can compare dissimilarity structures without regard to whether they are of the same dimension. For visualisation, only three compositional types are hypothesised, corresponding to three levels of darkness in each colour (e.g., blue, green, and red). Specifically, we hypothesised that dissimilarities between sites (or time points) of the same level of darkness were zero (i.e., identical composition), those between sites of one darker or lighter difference were 0.5, and those between the darkest and lightest sites were 1.0. The grey arrows indicate conventional dissimilarity (i.e., first-order dissimilarity) and the black arrows indicate HOD. Among compared datasets, black and white stars indicate identical sites or time points. The potential analytical framework for HOD based on spatial and temporal comparisons (i.e., spatial and temporal HOD) is shown in (b). The upper panel shows examples of spatial HOD, and the lower panel shows examples of temporal HOD. In both cases, the left-hand side shows examples of community structure comparisons between different guilds and intraspecific genetic structure between species, while the right-hand side shows examples of changes in spatial HOD or temporal HOD along a temporal or spatial axis.

### Higher-order dissimilarity

In biodiversity science, one dissimilarity value is generally calculated from two vectors of multivariate biological quantities (e.g., community composition or allele frequency). Specifically, a community dissimilarity index (e.g., Bray–Curtis dissimilarity; Odum 1950) and a genetic distance index (e.g., *F*_*st*_; Wright 1969) were calculated based on vectors of species abundance in two target communities and vectors of genetic allele frequency in two target populations, respectively. In addition, multiple vectors are summarised into a matrix called a site-species (or time-species) matrix (e.g., compositional vectors for sites or time points). The calculation of dissimilarity values for all pairs of vectors yields a dissimilarity matrix, which is a conventional pairwise dissimilarity; we call this first-order dissimilarity (Fig. 1a). Here, we extend the first-order dissimilarity to formally define HOD as dissimilarities of spatial or temporal dissimilarity structures (Fig. 1a). We also define ‘order’ as nested times for calculating dissimilarity, e.g., conventional pairwise dissimilarity is the first order; and dissimilarity of dissimilarity matrices the second order. By definition, the concept can be extended to a higher order where appropriate, such as to the third order (i.e., dissimilarity of second-order dissimilarity matrices).

The concept of HOD is a generalisation of the traditional analytical approach to identify species groups sharing the same trend and boundary in space and time (Holt et al. 2013; Gotelli et al. 2022). In addition, HOD targets the spatial structure, which has been the primary subject of biogeography and temporal structure. Specifically, to spatially and temporally distinguish HODs, we use the terms’ spatial HOD’ and ‘temporal HOD’ (Fig. 1b). Given the recent rapid growth of spatio-temporal monitoring data, HODs regarding temporal variation in spatial structure (i.e. temporal change in spatial HOD, Fig. 1b) or spatial variation in temporal structure (i.e. spatial change in temporal HOD, Fig. 1b) will be applicable in future analyses. The subsequent sections mainly focus on second-order dissimilarities similar to spatial and temporal HODs to elucidate the concept.

### Implementation of the higher-order dissimilarity

The most basic calculation of the HOD index (Fig. 2) is based on the RV coefficient (Escoufier 1973; Robert & Escoufier 1976), which is the multivariate generalisation of the squared Pearson correlation that correlates two matrices with corresponding sites (Blanchet et al. 2014). Before comparing two multivariate data matrices (i.e., conventional dissimilarity matrices), one may employ a dissimilarity index as a transformation technique for each data matrix. Specifically, principal coordinate analysis (PCoA) should be applied to each conventional dissimilarity matrix. This process yields rectangular data matrices, ensuring each dissimilarity matrix adheres to Euclidean properties. Then, the RV coefficient can be calculated through a comparison of two PCoA matrices (i.e., X and X’ in Fig. 2). The RV coefficient yields values ranging from 0 (indicating no correlation) to 1 (representing perfect correlation). Therefore, we compute the HOD index as one minus the RV coefficient to transform it into a dissimilarity measure, which we refer to as the contrast RV coefficient.

**Figure 2.**
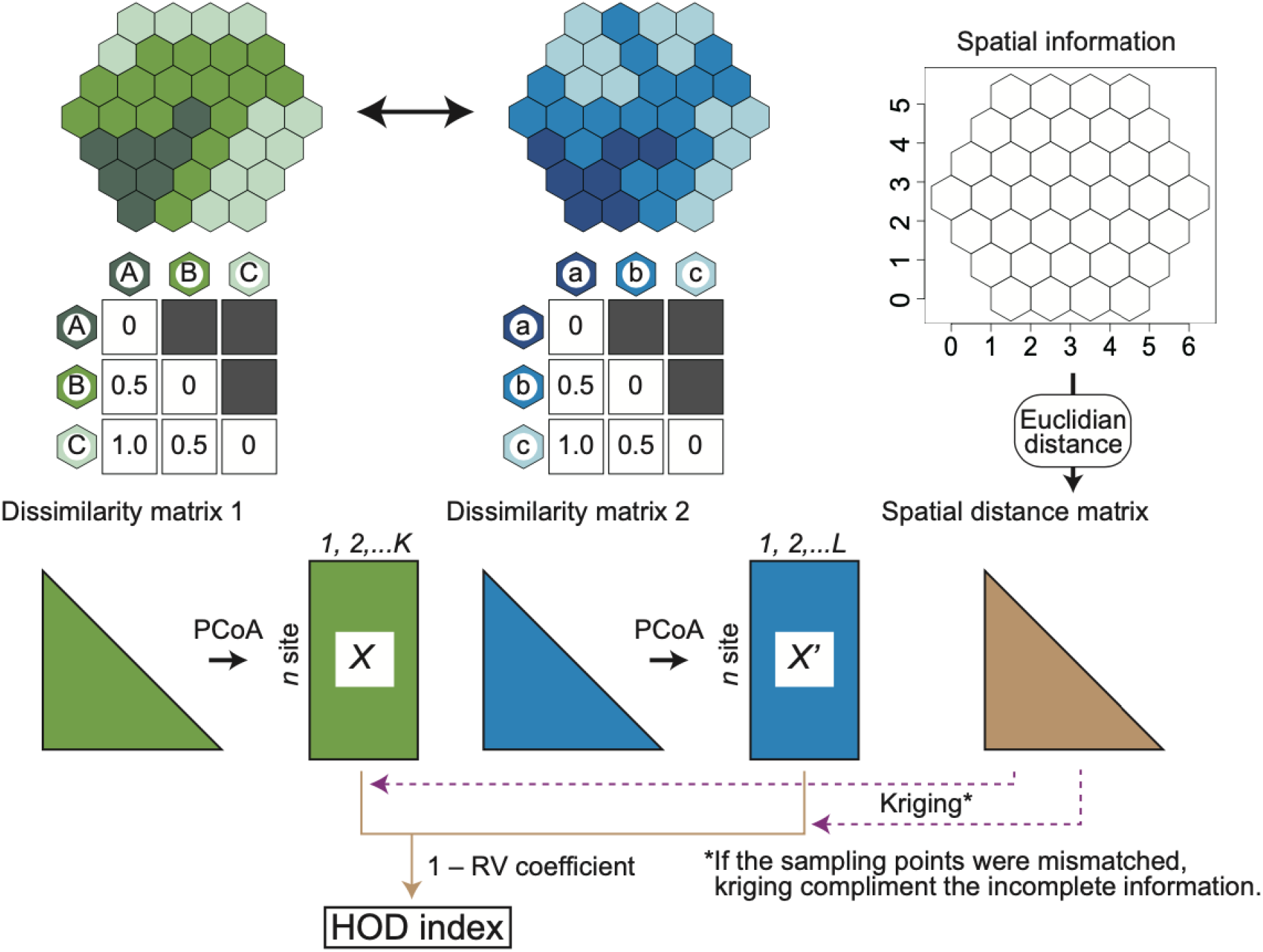
Schematic representation of specific analytical procedures for higher-order dissimilarity (HOD). For visualisation, only three compositional types are hypothesised, corresponding to three levels of darkness in each colour (e.g., blue, green, and red). Specifically, we hypothesised that dissimilarities between sites (or time points) of the same level of darkness were zero (i.e., identical composition), between sites of one darker or lighter difference were 0.5, and between darkest and lightest sites were 1.0. The asterisk notes the necessity of the Kriging process if the specific sampling points are not identical.

When sampling points comprising two biodiversity datasets are obtained at the same spatial or temporal location and correspond precisely one-to-one, their dissimilarity matrix elements also correspond. However, this is not always true. For example, when comparing two phylogeographic patterns, the sample points of each species are usually in different locations. In such a case, the elements of dissimilarity matrices do not correspond one-to-one and temporal or spatial proximity information is required for comparison. To calculate the HOD index for such an ‘irregular dataset’ (i.e., a dataset where the two compositional vectors to be compared were observed at different spatial and temporal points and did not correspond on a one-to-one basis), kriging, which is an interpolation method based on variograms (Wagner 2004; Legendre & Legendre 2012; Borcard et al. 2023), helps estimate the values of one data set at the locations of the other before the calculation. After interpolating the missing values, the methods for calculating the HOD index (i.e., contrast RV coefficient) remain the same as described above. However, spatio-temporal interpolation by kriging is not always effective and must be used cautiously when there is either no spatial/temporal autocorrelation or weak autocorrelation, or when there are disconnected patterns. One possible approach, based on spatial or temporal weight matrices such as Moran’s I (Wartenberg 1985; Lee 2001; Eckardt and Mateu 2021), could be effective except in the absence of complete spatial or temporal autocorrelation.

To check the properties of the HOD indices, we summarised the specific values of spatial HOD for pairs of simple sample cases in Table 1. While the degrees of spatial (or temporal) autocorrelation are not directly relevant to this issue in the case of complete datasets, they are included as reference information. When interpolating using kriging, the degree of autocorrelation would potentially affect the accuracy of the calculation. Here, spatial autocorrelation was obtained through regression analysis of distance-based RDA to the PCNM axes of spatial distances (Legendre et al. 2015). To calculate both the RV coefficient and the p-values using the function “coeffRV” of the *FactorMineR* package (Lê et al. 2008) in R (R Core Team, 2022).

**Table 1.**
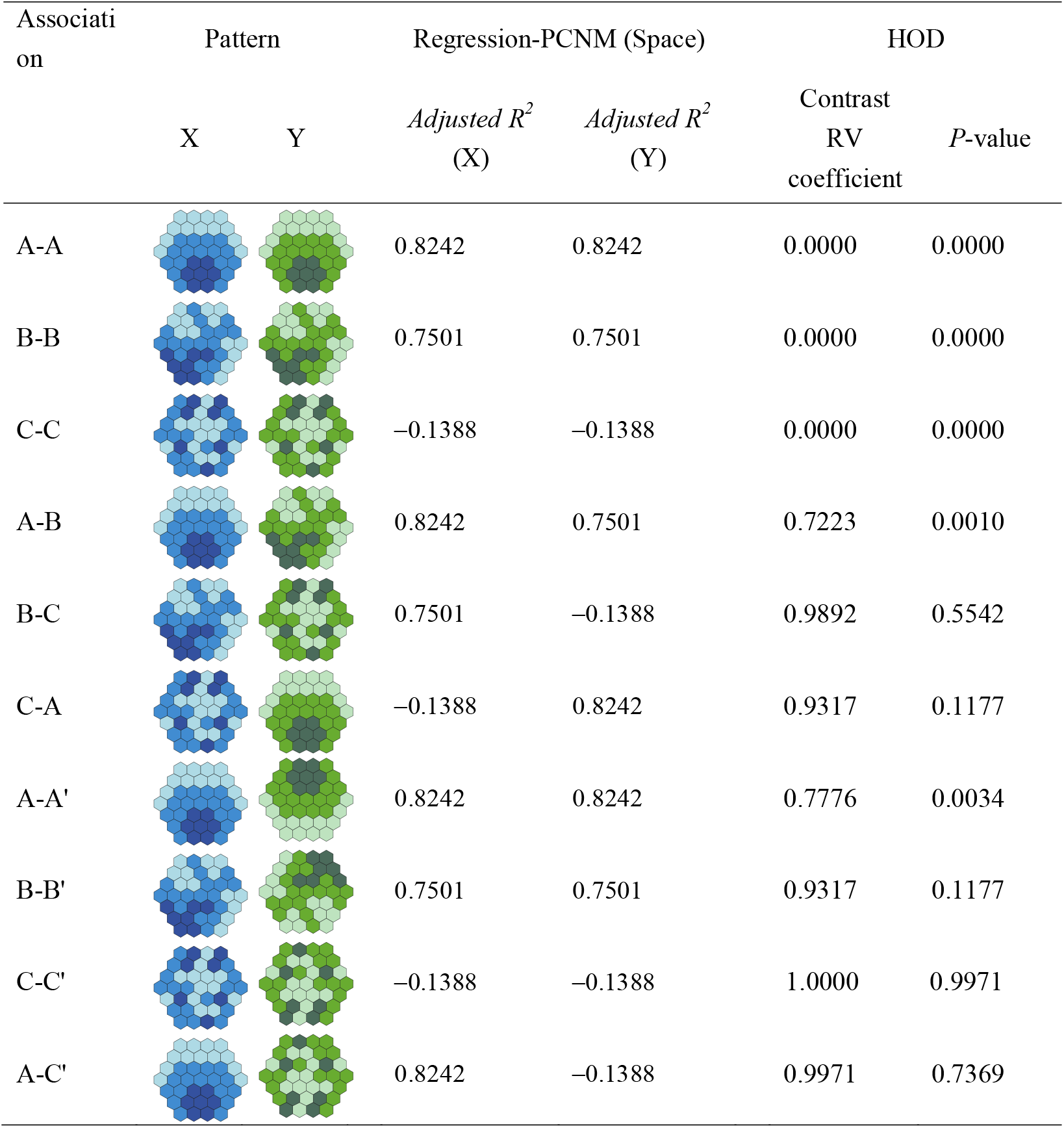
Simple sample cases to show properties of HOD indices. Patterns X and Y represent pairs for calculating HODs. *Adjusted R*^*2*^ (X) and (Y) indicate the degree of spatial autocorrelation for each pattern X and Y, obtained through regression analysis of distance-based RDA to the PCNM axes of spatial distances. The HOD indices calculated as contrast RV coefficient (i.e., 1 – RV coefficient) are shown. For visualisation purposes, only three compositional types are hypothesised, corresponding to three levels of darkness in each colour (e.g., blue and green). Specifically, we hypothesised that dissimilarities between sites of the same level of darkness were zero (i.e., identical composition), between sites of one darker or lighter difference were 0.5, and between the darkest and lightest sites were 1.0.

| Associati<br>on | Pattern |  | Regression-PCNM (Space) |  | HOD |  |
| --- | --- | --- | --- | --- | --- | --- |
|  | X | Y | *Adjusted R<sup>2</sup>*<br>(X) | *Adjusted R<sup>2</sup>*<br>(Y) | Contrast<br>RV<br>coefficient | *P*-value |
| A-A |  |  | 0.8242 | 0.8242 | 0.0000 | 0.0000 |
| B-B |  |  | 0.7501 | 0.7501 | 0.0000 | 0.0000 |
| C-C |  |  | -0.1388 | -0.1388 | 0.0000 | 0.0000 |
| A-B |  |  | 0.8242 | 0.7501 | 0.7223 | 0.0010 |
| B-C |  |  | 0.7501 | -0.1388 | 0.9892 | 0.5542 |
| C-A |  |  | -0.1388 | 0.8242 | 0.9317 | 0.1177 |
| A-A' |  |  | 0.8242 | 0.8242 | 0.7776 | 0.0034 |
| B-B' |  |  | 0.7501 | 0.7501 | 0.9317 | 0.1177 |
| C-C' |  |  | -0.1388 | -0.1388 | 1.0000 | 0.9971 |
| A-C' |  |  | 0.8242 | -0.1388 | 0.9971 | 0.7369 |

### Resolvable questions through the higher-order dissimilarity approaches

Theoretically, HOD concepts apply to any dissimilarity dataset if the relative position (i.e., sites or time points) can be determined based on the absolute distance between pairs of datasets. For example, the monitoring of the genetic structure is one of the most urgent targets of biodiversity monitoring (Hoban et al. 2021; O’Brien et al. 2022), although no effective monitoring framework has been developed. In this context, the HOD provides an ideal framework for the quantitative evaluation of genetic structure temporally (Fig. 3). As frequently probable examples, we show two hypothetical scenarios of genetic structure changes: one is a stable scenario (orange line and dots), and the other is a genetically homogenised scenario (blue line and dots) in Fig. 3 (see Supplementary Text 1 for detailed information about construction for visualisation). Fig. 3(a) indicates temporal changes in the values of the HOD index (i.e., contrast RV coefficient) to time 0 (starting point), and the lower Fig. 3(b) represents mapped overviews of changes in genetic structure over time, from which we calculated the HOD values. The scarcity of such spatially coordinated data on genetic structures over a wide area will soon be addressed, particularly following the development of environmental DNA technology (Tsuji et al. 2023), and our framework will be an essential contribution to genetic diversity monitoring.

**Figure 3.**
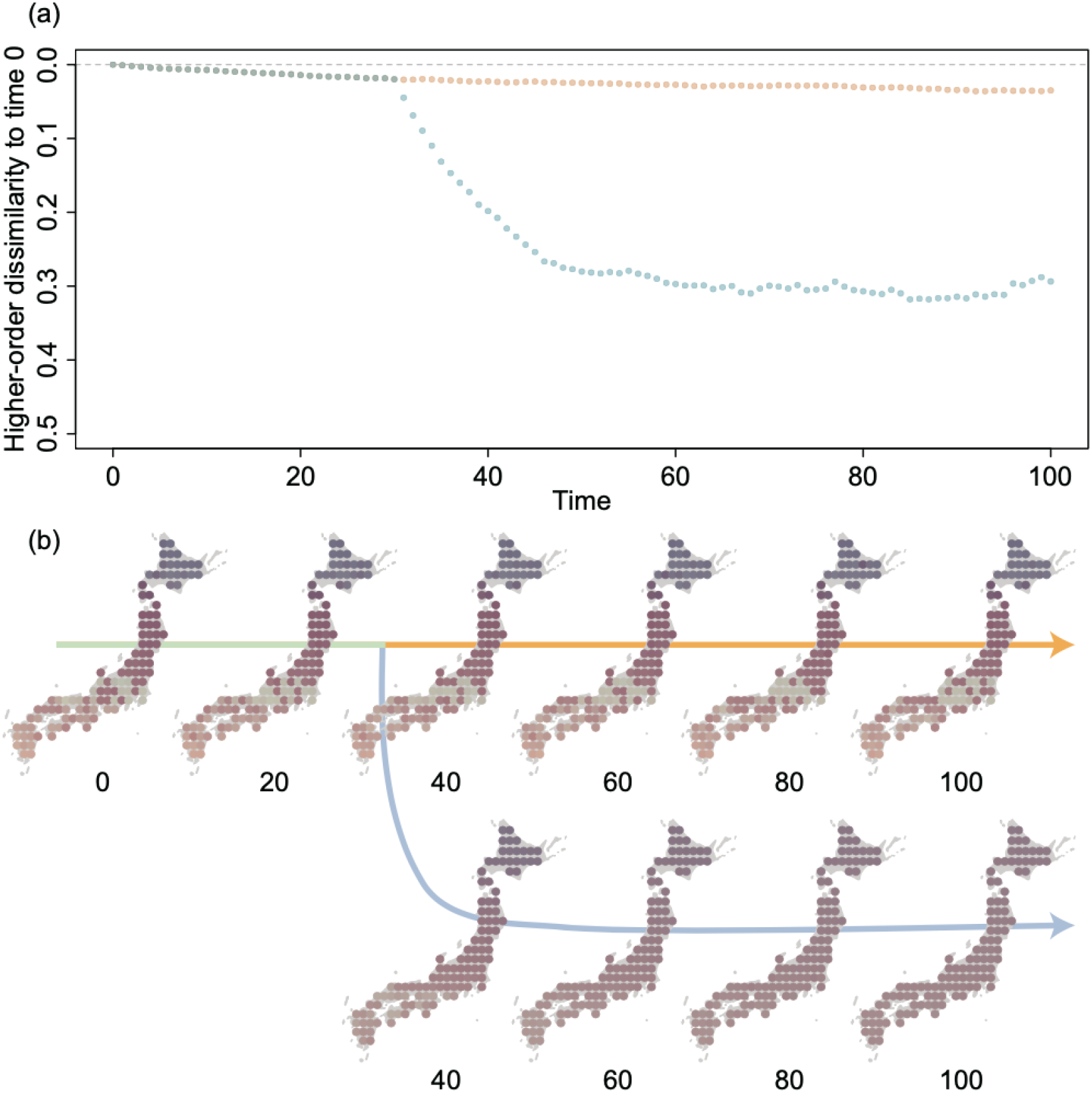
Sample application of higher-order dissimilarity (HOD) to genetic monitoring using two simulated datasets based on two types of scenarios from time 0 to time 100. In both scenarios, the stable conditions are hypothesised by time 30 (shared section in light green). After time 30, the genetic structure is stable in one scenario (upper section in orange) but homogenised in the other (lower section in blue). The change of the calculated HOD index (i.e., contrast RV coefficient) compared to the value at time 0 is shown in (a). The genetic structure is visualised in (b), and the colours were determined using the two axes of NMDS (details see Supplementary text 1).

To show the advantages of HOD more comprehensively, we summarised the potential applications of HOD in Table 2, including five types of datasets: community composition, genetic frequency, ecosystem function, interaction network, and phylogeny. Most cases are straightforward extensions of those shown in Fig. 1b. One particular case is that HOD approaches are applicable to phylogenetic patterns despite the difficulty in extracting single-dimensional values, for example, the output of non-metric multidimensional scaling based on multiple traits. HOD approaches can be applied to various community ecology, biogeography, phylogeography, and macroevolution datasets. Particularly in the case of examples covering macroevolution (i.e., phylogenetic structures), a previous study had a similar, though not identical, perspective (Duarte 2011). This fact further emphasises the importance of developing a field of study for HOD.

**Table 2.**
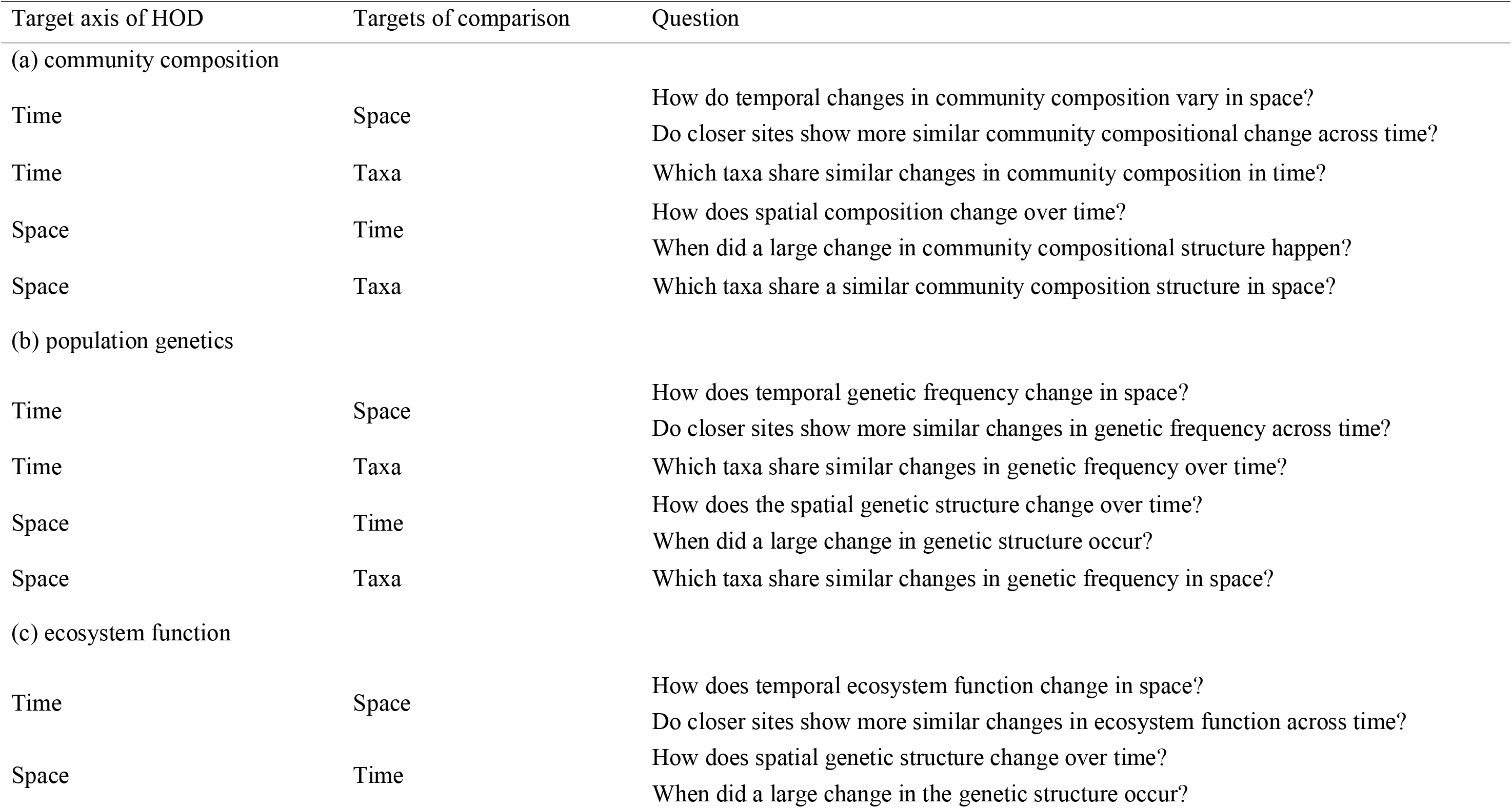
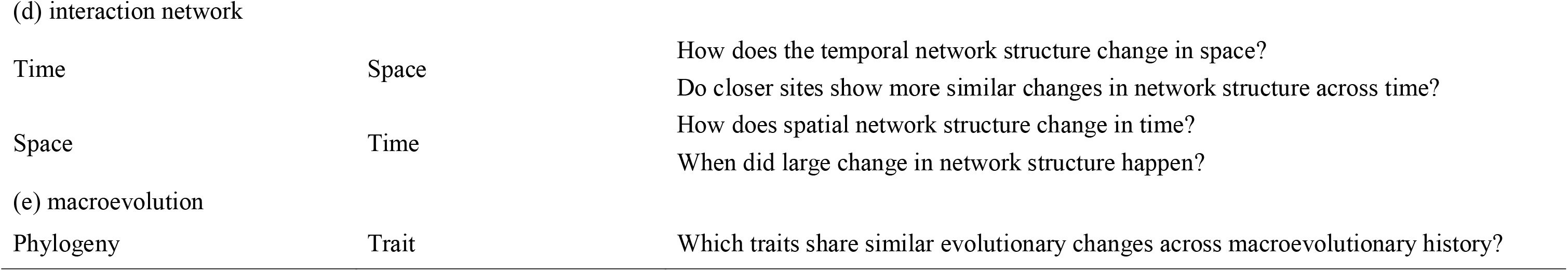
Potential applications of the higher-order dissimilarity (HOD) approach.

| Target axis of HOD | Targets of comparison | Question |
| --- | --- | --- |
| (a) community composition |  |  |
| Time | Space | How do temporal changes in community composition vary in space? |
|  |  | Do closer sites show more similar community compositional change across time? |
| Time | Taxa | Which taxa share similar changes in community composition in time? |
| Space | Time | How does spatial composition change over time? |
|  |  | When did a large change in community compositional structure happen? |
| Space | Taxa | Which taxa share a similar community composition structure in space? |
| (b) population genetics |  |  |
| Time | Space | How does temporal genetic frequency change in space? |
|  |  | Do closer sites show more similar changes in genetic frequency across time? |
| Time | Taxa | Which taxa share similar changes in genetic frequency over time? |
| Space | Time | How does the spatial genetic structure change over time? |
|  |  | When did a large change in genetic structure occur? |
| Space | Taxa | Which taxa share similar changes in genetic frequency in space? |
| (c) ecosystem function |  |  |
| Time | Space | How does temporal ecosystem function change in space? |
|  |  | Do closer sites show more similar changes in ecosystem function across time? |
| Space | Time | How does spatial genetic structure change over time? |
|  |  | When did a large change in the genetic structure occur? |
| (d) interaction network |  |  |
| Time | Space | How does the temporal network structure change in space?<br>Do closer sites show more similar changes in network structure across time? |
| Space | Time | How does spatial network structure change in time?<br>When did large change in network structure happen? |
| (e) macroevolution |  |  |
| Phylogeny | Trait | Which traits share similar evolutionary changes across macroevolutionary history? |

The main focus has been on comparisons between dissimilarity matrices with similar properties, but this is not necessarily true. Examples include comparisons among communities of different trophic levels and comparisons between community structures and genetic structures in community members. In traditional approaches, revealing the relative importance of the driving processes of local communities, such as abiotic vs. biotic factors and/or top-down vs. bottom-up processes as drivers of community structures, has been a major concern (Smith et al. 2010; HilleRisLambers et al. 2012). The HOD approach is a different perspective that directly evaluates the similarity of patterns without explicitly considering the driving processes. Low values of HOD among data from different trophic levels of communities and different hierarchies of genetic and community data allowed us to generate novel hypotheses with more direct evidence, as the processes shaping them are considered common among the compared targets (taxonomic differences; Fig. 1b). Conversely, high HOD values suggest that the process of constructing the overall system is complex and contingent. From an application perspective, high values of spatial HOD also suggest that achieving conservation through establishing protected areas, setting focal species for conservation, or implementing a single conservation policy would be difficult, as the driving processes differ among taxa and levels of biological entities.

### Future challenges

In the HOD approaches proposed here, we cannot distinguish between parts of continuous patterns and, indeed, disjunctive patterns in dissimilarities, for example, either simple distance-decay patterns or patterns due to isolation by geographic barriers. Therefore, where the commonality of spatial disconnections is the main concern, combining HOD approaches with existing approaches would be more effective in identifying disjunct barriers (e.g., Soltis et al. 2006). In temporal structures, approaches separating trend components have been used (Cowpertwait & Metcalife 2009), and potentially combined with such methods, it may be possible to evaluate continuous and disjunctive dissimilarities separately. Furthermore, spatially- or temporally-constrained clustering approaches (Guénard & Legendre 2022) based on the HOD matrix can depict disjunctive patterns of higher-order structures in community compositions. However, further testing is required to clarify how much complex hierarchy is allowed for a sufficiently accurate evaluation for calculating the HOD index.

Although we mainly focused on the second-order dissimilarities in the present study, the applicability of third- and higher-order dissimilarities would increase when the amount and dimension of biodiversity information continue to grow. To illustrate the benefit of HOD, we showed temporal changes in genetic structure (Fig. 3). The figure shows a pattern in a single species and, thus, second-order dissimilarity (i.e., space × time). If this dataset is available for multiple species, the similarity and dissimilarity of temporal changes in spatial structure in population genetics can be quantitatively evaluated among species using third-order dissimilarities (i.e., space×time×species). In such multidimensional situations, a hierarchical multiple-factor analysis (Le Dien & Pagès 2003) can be a useful exploratory tool for grasping the structure of communities that affect the higher-order HOD. This simple example tells us that the breadth of areas that the HOD concept can adopt is rapidly expanding, aligning with the forthcoming explosive increase in available dissimilarity information.

### Conclusions

The rapid increase in the availability of spatial and temporal biodiversity information necessitates comparative studies on spatio-temporal structures of biodiversity compositional patterns, which require further development of the dissimilarity concept over conventional ones to consider higher dimensions of dissimilarity: dissimilarity of dissimilarity structures. Our study introduced the concept of higher-order dissimilarity (HOD), which can account for dissimilarities of spatial or temporal dissimilarity structures. For example, this framework can be applied to various types of broad-scale biodiversity monitoring datasets and enables the evaluation of temporal changes in the spatial HOD to a baseline. Furthermore, the concept is applicable even if the compared datasets have entirely different origins (e.g., different taxonomic groups and hierarchies of biological organisation, Fig. 1b) as long as they are summarised into dissimilarity information at their level. We mainly focused on the biodiversity dataset in the present study; the concept of HOD applies to all kinds of dissimilarity matrices, not just those based on biodiversity. For example, various contemporary issues arise in the social-ecological system, and this HOD will be useful in examining the relationships between different layers, such as the economy, culture, and biodiversity. Therefore, the HOD concept and related approaches pave the way for novel dimensional approaches that deal with large amounts of dissimilarity information.

## Acknowledgements

We thank Dr. Satoshi Aoki for his advice on choosing a coalescent simulator and Prof. Pierre Legendre for his insightful comments on the manuscript, drawing from his profound and expert knowledge.

## Data availability

Simulated data and codes will be uploaded after acceptance of the manuscript to Figshare.

## Author contributions

RN, FI, KF, and TK designed the study; RN conceived the idea of higher-order dissimilarity, analysed the datasets, and wrote the first draft of the manuscript, with significant inputs from KF, FI, and TK; KF and RN developed the methodology to quantitatively evaluate the idea introduced in the present study; and FI conceived the basic concept of the methodology. All authors have contributed substantially to the final version of this manuscript.

## Appendices

**Figure S1** Hexagon grids used in simulation.

**Supplementary text 1** Detailed information on simulated genetic structure across four major islands in Japan.

