## Supplementary figures and images for "Higher-order dissimilarity in biodiversity: Identifying dissimilarities of spatial or temporal dissimilarity structures"

### Figure S1

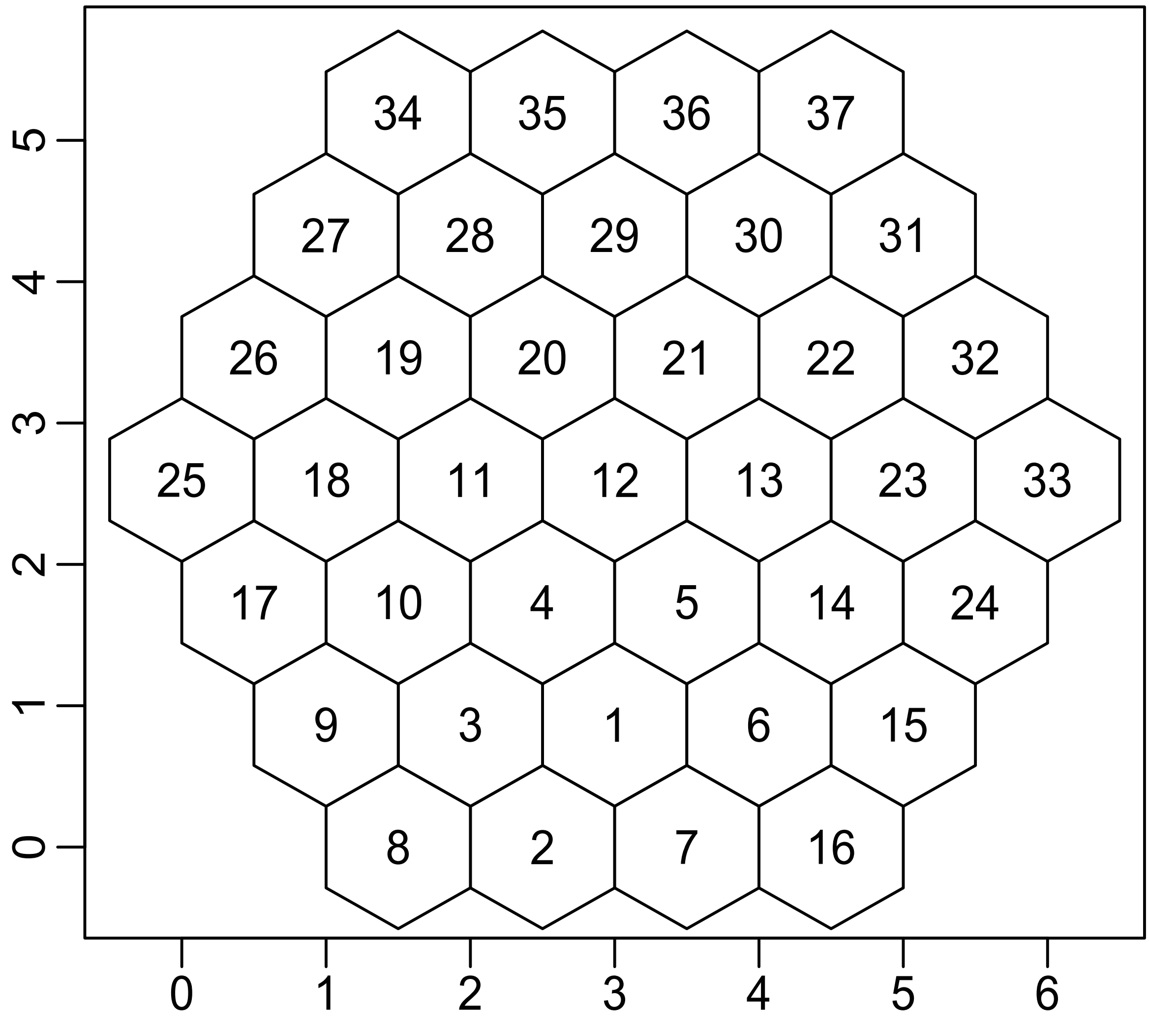
