## Supplementary text for "Higher-order dissimilarity in biodiversity: Identifying dissimilarities of spatial or temporal dissimilarity structures"

**Supplementary text 1.** Detailed information about simulated genetic structure across four major islands in Japan

A coalescent simulation was conducted using the software *ms* (Hudson 2002) to construct the simulated datasets of the genetic structure as a sample case. A modified version of *ms* (*msutilis*; <https://github.com/heavywatal/msutils> accessed on 2021.12.12) was used in the present study. The modified software used the dSFMT (ver. 2.2.3; Matsumoto & Nishimura 1998), which is a successor to the Mersenne twister instead of a linear congruential generator in the original *ms* to generate the pseudorandom number because the initially implemented pseudorandom number generator yielded inferior performance (Park and Miller). To facilitate the provision of specific images for empirical monitoring studies, we conducted a genetic simulation of Japanese archipelagoes as an example. First, we determined 113 hypothetical populations on the four major islands using the function "*population"* in the package *slendr*.

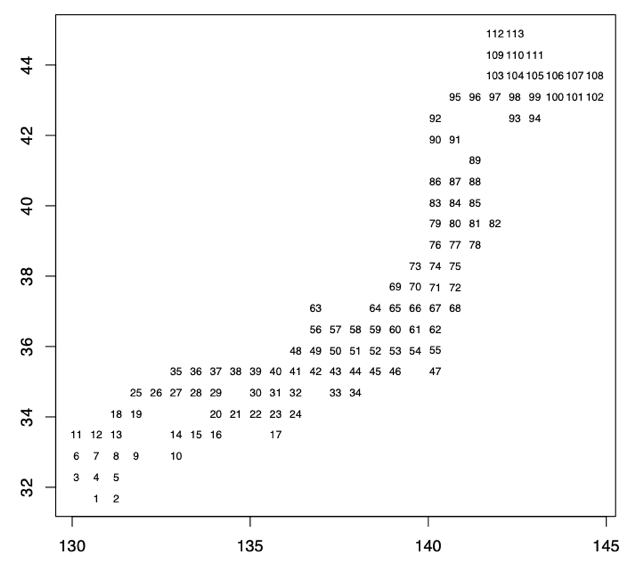

We hypothesised that gene flows only occur among adjacent populations at a fixed rate (2.5), and gene flows between populations isolated by the sea are set to be relatively lower (0.5) than those in the *msutilis* simulation. We hypothesised that diploids without chromosomal recombination would simplify the simulation. As outputs, we set 50 individuals with 10 chromosomes for each of the 113 populations and 10 numbers (e.g. 0001010101) for each chromosome. The ancestral and derived states were coded as 0 and 1, respectively. For each chromosome, 10 numbers were converted from binary (e.g. 0001010101) to decimal (e.g. 341). After the conversion, they were clustered into groups of four digits for alignment, and 9 were added to the first line to be considered as a single microsatellite marker on a single chromosome, that is, genotypes for each chromosome (e.g. 90341) to determine individual genotypes. We further simulated two types of genetic homogenisation scenarios based on the simulated results of population genetics. Specifically, we simplified homogenisation as the movement of chromosomes among populations and changed the degree of homogenisation between the two scenarios. Each chromosome is capable of leaving 0 to 3 chromosomes of the same type. Furthermore, the chromosomes are dispersed to other populations with a certain probability (i.e. homogenisation) depending on the distance, as mentioned above. Among the chromosomes in a population, 100 were finally selected and passed on to the next generation. The scenario of high homogenisation was hypothesised to be four times higher than that of low homogenisation in the later stage of the simulation. The two scenarios share the same hypothetical condition: low genetic homogenisation until 30-time steps. In one scenario (i.e., the upper scenario in Fig. 3ab [orange circles and orange line]), the condition of low genetic homogenisation continued until the end of the simulation. In the other scenario (i.e. lower scenario in Fig. 3ab [blue circles and a blue line]), higher genetic homogenisation was hypothesised from 31-time steps to the end (i.e., 100-time steps). All pairs of genetic differentiations in each simulation were calculated based on Da (Nei 1987) using the "*genet.dist*" function in *hierfstat* package (Goudet 2005). We further calculated the HOD type 1 value between the dissimilarity pattern at time 0 (i.e. the starting point) and that of all other time steps in each simulation. The colours of each population in Fig. 3 were determined based on the two values along the two measurement axes of non-metric multidimensional scaling. Non-metric multidimensional scaling was calculated using the "*metaMDS*" function in the *vegan* package (Braak et al. 2022). All analyses, except the simulation for population genetics, were conducted in R (ver. 4.10; R core team 2021).
